# Weighted transitivity scores that account for triadic edge similarity in undirected graphs

**DOI:** 10.1101/2022.01.11.475816

**Authors:** Guillaume Peron

## Abstract

The graph transitivity measures the probability that adjacent vertices in a network are interconnected, thus revealing the existence of tightly connected neighborhoods playing a role in information and pathogen circulation. When the connections vary in strength, focusing on whether connections exist or not can be reductive. I score the weighted transitivity according to the similarity between the weights of the three possible links in each triad. I illustrate the biological relevance of that information with two reanalyses of animal contact networks. In the rhesus macaque *Macaca mulatta*, a species in which kin relationships strongly predict social relationships, the new metrics revealed striking similarities in the configuration of grooming networks in captive and free-ranging groups, but only as long as the matrilines were preserved. In the barnacle goose *Branta leucopsis*, in an experiment designed to test the long-term effect of the goslings’ social environment, the new metrics uncovered an excess of weak triplets closed by strong links in males compared to females, and consistent with the triadic process underlying goose dominance relationships.

## Introduction

Social interactions between individuals are rarely binary. Instead, the connections between individuals vary in strength or valence. And even if the instant interactions indeed have a binary outcome, the consequences usually accumulate over several interactions, e.g., the fitness benefits of long-lasting bonds (Silk et al., 2009), and the hierarchy emerging from successive agonistic interactions (Franz et al., 2015). In that context, it seems useful to develop network summary metrics that account for edge weights (Barrat et al., 2004), instead of dichotomizing the network or to scale the timeframe down until interactions can be considered binary. Here I focus on the graph transitivity (Newman et al., 2002). The graph transitivity answers the question: when A is separately connected to B and to C, what is the probability that B and C are also connected. The transitivity influences the occurrence of indirect interactions (when B and C are not connected) as well as the expected shortest path length influencing the circulation of pathogens and information.

In its usual definition for binary networks, the graph transitivity corresponds to the relative frequency of closed and open triplets (Newman, Watts, & Strogatz, 2002; see also Watts & Strogatz (1998) and Barrat et al. (2004) for different definitions that use vertex-level scores instead of triplet enumeration).

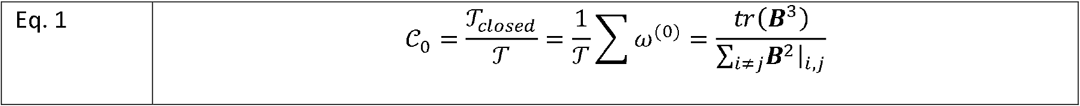

The sum in the middle is over triplets, i.e., any three individuals with at least two non-null links between them. 𝒯 is the total number of triplets in the network 𝒯_*closed*_, is the number of closed triplets among these. ω^(0)^ equals 1 if the triplet is closed and 0 if the triplet is open. Note that each closed triangle (*a, b, c*) contributes three closed triplets (*a, b*), (*b, c*), and (*a, c*) (Fig. 1). The last term represents a matrix formulation with ***B*** the binary adjacency matrix featuring a 1 in cell (*i,j*) if *i* and *j* are connected, and 0 otherwise.

**Fig. 1:**
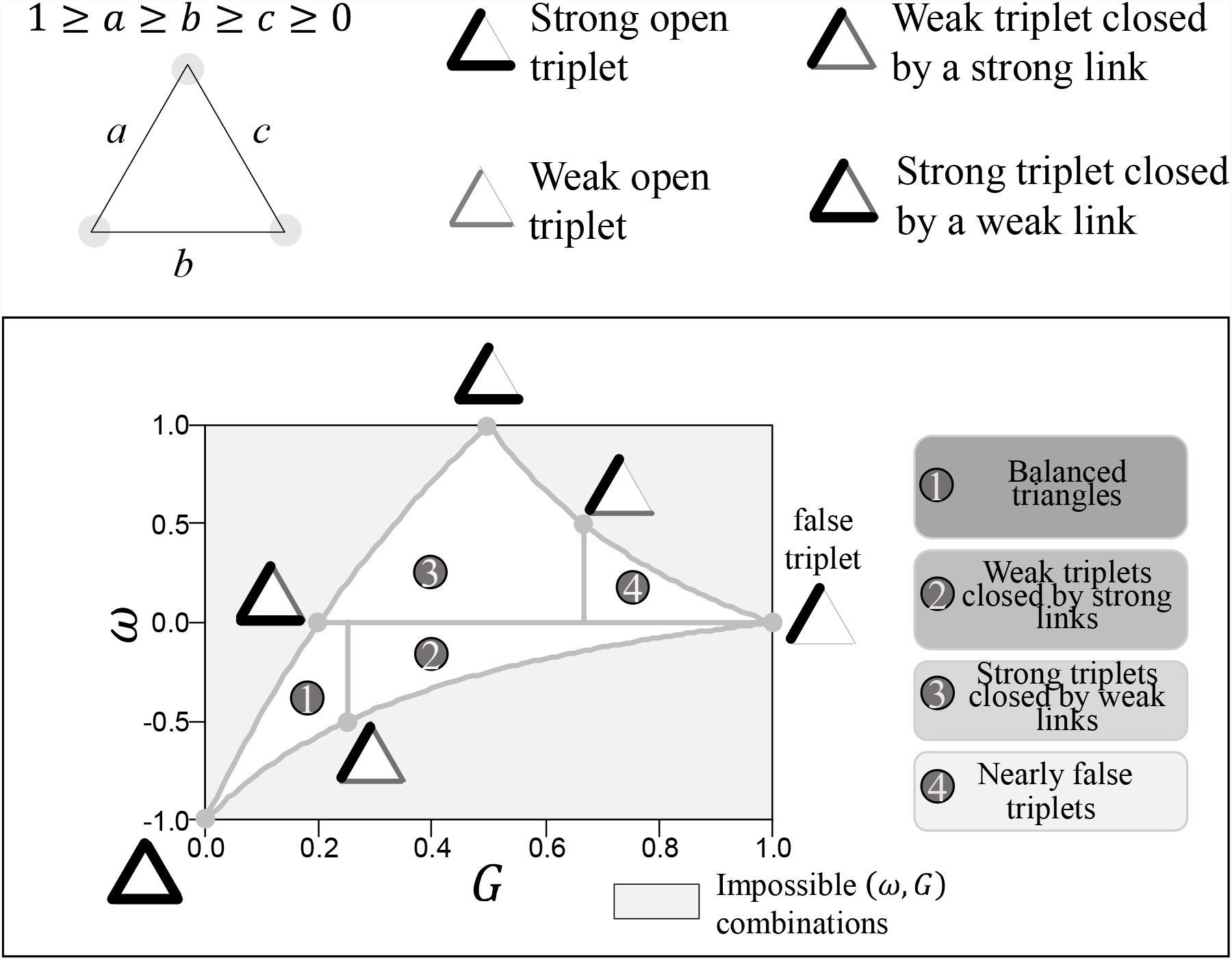
Triadic edge weight similarity and the four quadrants.

In weighted networks, the distribution of edge weights can significantly modify the transitive properties. For example, a triangle that features two strong links and one weak link may functionally operate more like an open triplet. Inversely, weak links that would maybe be considered insignificant when dichotomizing the network for transitivity evaluation may actually perform a significant role.

There already exists a weighted version of 𝒞_0_, derived by Opsahl and Panzarasa (2009). They use the geometric mean of the two links in each triplet as the weight of that triplet: 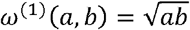 (or alternatively, the arithmetic mean, or the minimum, or the maximum). Then they derive the weighed transitivity as follows.

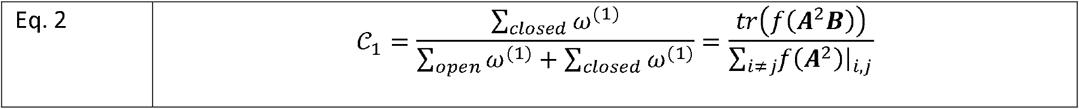

The second term represents a matrix formulation where ***A*** is the weighted adjacency matrix featuring the strength of the link between *i* and *j* in cell (*i,j*), ***B*** is the dichotomized version of ***A***, and *f* is a function that takes the square root of each cell.

Eq. 2 is attractive because 𝒞_1_ verifies most of the properties of 𝒞_0_ (Opsahl and Panzarasa, 2009). The main use of 𝒞_1_ is to assess the effect of the triplet weight on the probability that the third link in the triplet is created (Opsahl and Panzarasa, 2009). If 𝒞_1_ > 𝒞_0_, strong triplets (with a high ω^(1)^) are more frequently closed than weak triplets (with a low ω^(1)^). Inversely, if 𝒞_1_ < 𝒞_0_, then weak triplets are more frequently closed than strong triplets.

My grievances with Eq 2 are: (*i*) Eq. 2 does not capture the information about the similarity between the links: *a* and *b* can be very different or very similar, and yet yield the same ω^(1)^ value. (*ii*) Eq. 2 still involves some dichotomization: the strength of the closing link does not influence the triplet weight (Barrat et al., 2004). The only way to assess the effect of the weight of the third link, in an ad hoc way, would be to change the threshold *c*\* that is used to dichotomize the closing link. In addition to being relatively cumbersome and hard to interpret biologically, this procedure is expectedly unstable owing to the decrease in sample size when *c*\* increases. It also introduces an artificial nonlinearity at the threshold. Moreover, there rarely exists enough information to decide on a biologically meaningful threshold. The choice of *c*\* is often arbitrary (e.g., “more than one contact”, “statistically significant association”).

I propose new scores designed to capture the information about the similarity between the three edge weights in each triad, without any dichotomization required. I replaced the triplet weights ω^(0)^ and ω^(1)^ by a measure of the imbalance between the three links *a, b*, and *c*. I identified two ways to do so. The first is the rescaled sum of proportional pairwise differences:

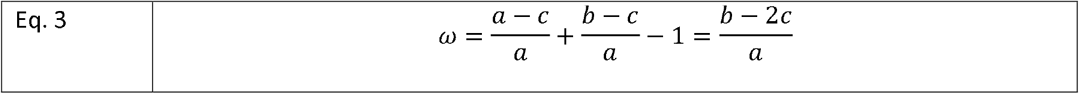

The rationale is that w compares *c* to *a* and to *b*. ω= 0 for a false triplet (*b* = *c* = 0), ω= 1 for an open triplet made of two equally strong links, and ω= -1 for a closed triangle made of three equally strong links (Fig. 1). The second option is the Gini coefficient:

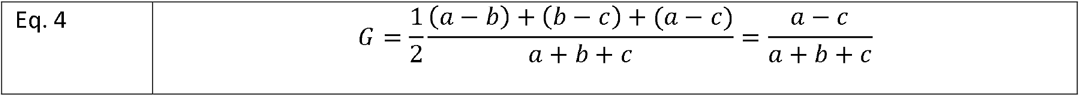

The rationale is that the Gini coefficient is a classical measure of inequality, expected to capture the three-way relationship between *a, b* and *c*. *G*= 1 for a false triplet (*b* = *c* = 0), *G*= 0.5 for an open triplet made of two equally strong links, and *G*= 0 for a closed triangle made of three equally strong links (Fig. 1). Note that throughout, the strengths of the links *a, b*, and *c* are rescaled to vary between 0 (no link) and 1 (strongest recorded link).

There is covariation between *ω* and *G*, and in particular, owing to the constraint that 1 ≥ *a* ≥ *b* ≥ *c* ≥0, the domain of possible combinations (*ω,G*) is restricted (Fig. 1). Despite this covariation, the two metrics are complementary and need to be interpreted together. For example, *ω* tends towards zero either when the triad is a false triplet dominated by a single link, or when the triad is a closed triangle but one of the links is about half as strong as the other two (Fig. 1). The value of *G* separates between the two situations (Fig. 1).

In other words, the aim of the method is to reduce the 3D space {*a, b, c* | 1 ≥ *a* ≥ *b* ≥ *c* ≥ 0) into a 2D space (*w, c* | 1 ≥ *a* ≥ *b* ≥ *c* ≥ 0), for inference about triadic association patterns.

For interpretation purposes, I suggest to divide the {*ω,G*} space into four quadrants (Fig. 1). I devised a permutation procedure, hereafter termed the quadrant test (Appendix S1), to assess whether the observed distribution of triads in the {*ω,G*} space depends only on the dyadic edge weights, or also depends on triadic association patterns. When the quadrant test is negative, it means that most of the observed variation in (*ω,G*) comes from the dyadic edge weights. When it is positive, triadic associations are biased towards specific combinations of edge weights.

Importantly, this permutation procedure removes all triadic processes. This would include automatic associations based on the finitude of shared resources. More precisely, when the edge weights are computed based on the proportion of time spent in close proximity, or on the proportion of a budget spent on joint ventures, transitivity may arise mechanistically. If individual A spends most of its time with B and C, then mechanistically B and C also spend most of their time together, even if they were here for A not for each other (Granovetter, 1973). Therefore, the interpretation of the quadrant test as formulated above critically depends on the way the edge weights are computed and also on the realized, empirical distribution of dyadic edge weights.

Next, one can compute graph-level transitivity scores based on *ω* and *G*. They are simply the average of *ω* and *G* over all the triads.

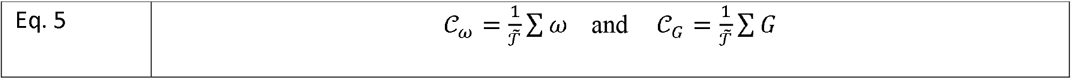

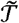 is the number of triads (a triad being any three individuals with at least one non-null link between them, counting one triad per triangle). Note that in Eq. 1 and 2 the sum was over triplets (a triplet being any three individuals with at least *two* non-null links between them, counting three triplets per triangle). The reason for this change is that the missing edges are now considered as an edge with weight zero. Therefore, despite the parallelism of form, 𝒞_*ω*_ and 𝒞_*G*_ do not compare directly to 𝒞_0_ or 𝒞_1_.

To assess the null hypothesis that the observed transitivity scores are expected from the edge density and edge weight distribution, I devised another permutation procedure, hereafter termed the distance test (Appendix S1). This test is similar in philosophy to the comparison between the usual transitivity score 𝒞_0_ and the edge density 𝒟_0_. I used the Erdős–Rényi model to generate a large number of random networks with the same number of vertices, same number of edges, and the same distribution of edge weights as the focal network (routine `sample_gnm` in `igraph`; Csardi & Nepusz, 2006). Next I computed the Mahalanobis distance between the observed (𝒞_*ω*_, 𝒞_*G*)_ value and the distribution of (𝒞_*ω*_, 𝒞_*G*_) among the simulated graphs. I computed the p-value of that distance under a chi-squared test with 2 degrees of freedom. If the test is positive, the observed transitivity scores are statistically different from the expectation at random. Note that extreme values of (𝒞_*ω*_, 𝒞_*G*_), in the lower left or lower right corners of the domain of definition, tend to test positive because the randomization is constrained in these corners. In addition, like for the quadrant test, the interpretation of the distance test critically depends on the way the edge weights are computed, and in particular whether the edge weights involve the expenditure of a finite shared resource like time or money (cf. quadrant test above).

In the following, I report on a simulation study and on two case studies to demonstrate the biological relevance of the new scoring method.

## Material and methods

### Simulation study

The envisioned main use of the new method is to diagnose whether a dense graph exhibits an excess of balanced triangles, of weak triplets closed by strong links, or of strong triplets closed by weak links. To illustrate that this task is achieved, I simulated transitive networks with those exact properties.

Scenario 1: excess of balanced triangles. I generated random full networks in which, at random, half of the edges were assigned a weight of 1 and half of the edges were assigned a weight drawn at random between 0 and 0.5.

Scenario 2: excess of strong triplets closed by weak links. First I generated random uniform networks with a c_O_ score of 0.8, using the rguman routine in R-package sna (Butts, 2020). Next, in each triangle, I assigned to one of the three links a low weight, drawn at random between 0 and 0.1, whereas the other two links in the triangle still had a large weight of one.

Scenario 3: excess of weak triplets closed by strong links. I generated random full networks in which, at random, 85% of the edges were assigned a weight of 0.1 and 15% of the edges were assigned a weight drawn at random between 0.75 and 1.

In addition to the three scenarios above, I considered a range of usual network types just to explore what their scores would be and whether they would be different from Scenarios 1-3: full networks with uniform or Gaussian distribution of edge weights, half-full networks in which I removed half of the edges at random, lattice graphs, modular graphs, star graphs, and scale-free graphs (Fig. 2). All these were generated using `igraph` for R (Csardi and Nepusz, 2006).

**Fig. 2:**
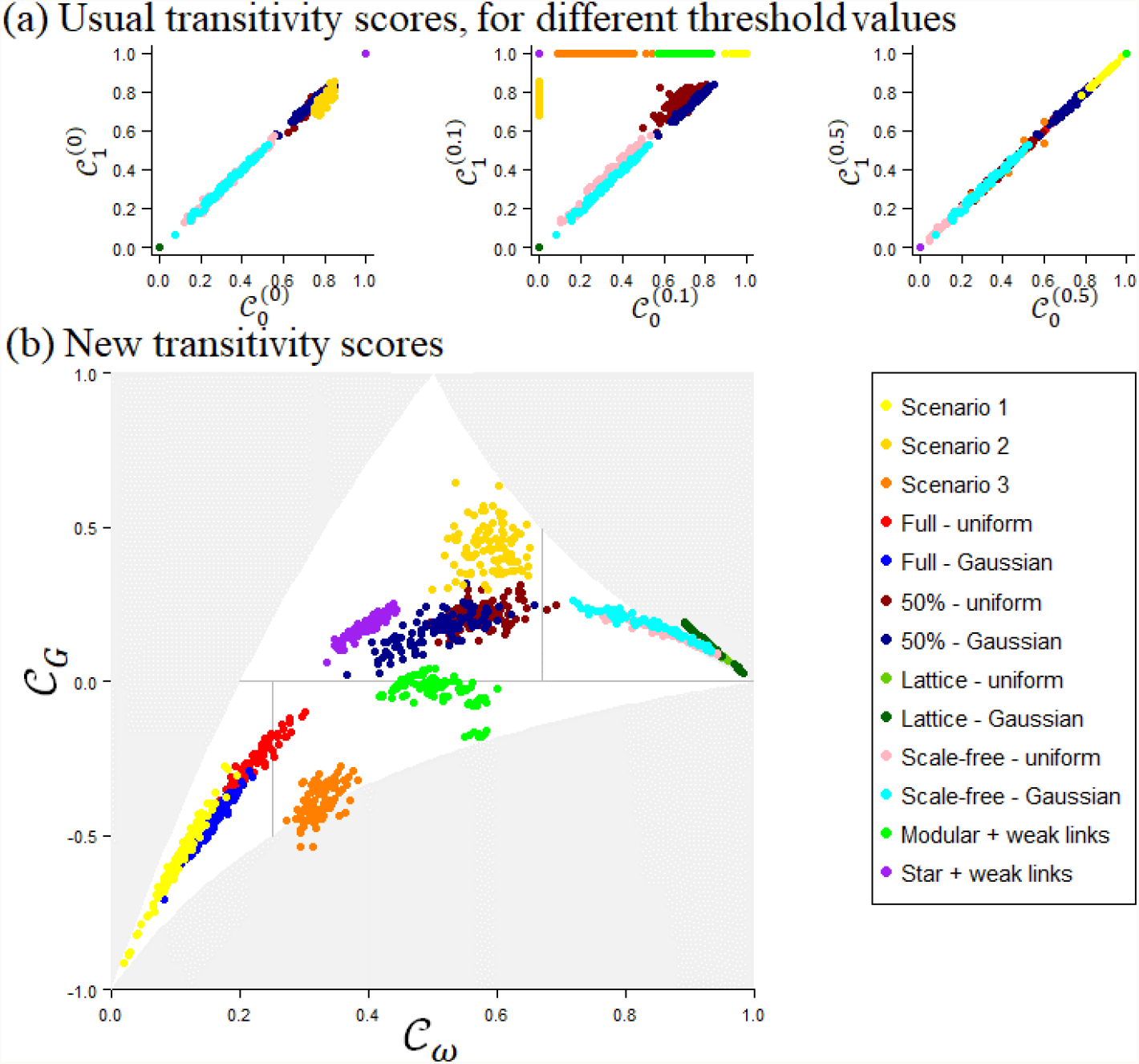
Simulation results. (a) The usual transitivity scores, with notation as in the main text; the superscripts indicate the value of the threshold *c*\* to decide when a link is significant. (b) The new transitivity scores. Scenarios 1-3 as explained in the main text. “full” = all the vertices are connected to each other but the strength of the connection varies according to a uniform or Gaussian law. “50%” = same as “full” but half of the edges are removed. “Lattice”: graph with a regular tiling pattern creating short-range neighborhoods for each vertex. I included lattices with 1 dimension (chain), 2 (square grid), 3 (cubic grid), and 4 dimensions. “Scale-free”: networks whose degree distribution follows a power law, meaning that the vertex with the most connections has exponentially more connections than the vertex with the second most connections, and so on. “Modular + weak links”: I created 2 to 8 full graphs with strong within-graph connections, then connected them to each other with edge weights below 0.1. “Star + weak link”: all the vertices were connected to a central vertex with strong links, and then among themselves with edge weights below 0.1.

#### Illustration 1: grooming networks in rhesus macaques

Rhesus macaques (*Macaca mulatta*) are a social primate typically found in multimale groups that can range in the hundreds of individuals. They exhibit an “intolerant social style” characterized by steep dominance gradients, frequent displays of aggression and submission, and, importantly, strong preference for kin in social interactions (Thierry, 2007). Due to this social style, my working hypothesis is that the kinship structure of the group, i.e., whether the matrilines are preserved or not, influences the emerging network properties. I reanalayzed 6 datasets of grooming interactions within groups of rhesus macaques with or without matrilines. As a control, I also tested the effect of captivity vs. free-ranging. Two free-ranging groups, both from the same locality but different time periods, featured the naturally-occurring matrilines (Griffin and Nunn, 2012; Puga-Gonzalez et al., 2018). One captive group featured locally born adult and immature offspring, therefore similar to the natural kinship structure (Massen and Sterck, 2013). In the last three groups, all from the same study in captivity, the individuals came from various origins and met each other after weaning age (Balasubramaniam et al., 2018). Edge weights corresponded to the frequency of recorded allogrooming interactions for each dyad during standardized observation sessions. I did not distinguish who was the groomer and who was the recipient, i.e., symmetric, undirected edges.

#### Illustration 2: barnacle geese experiment

Barnacle geese (*Branta leucopsis*), like all geese, are among the most gregarious of vertebrates, yet exhibit high levels of competition for food and frequent agonistic behavior (Black and Owen, 1989; Stahl et al., 2001). Within geese flocks, kin and mate support is essential to assert dominance (Black and Owen, 1989; Kurvers et al., 2013). I reanalyzed sub-group memberships in the captive population that Kurvers et al. (2013) maintained in an experimental environment. The edge weights corresponded to the frequency of standardized sampling occasions during which two individuals occurred together on the same feeding patch, as opposed to foraging in separate sub-groups.

My working hypothesis is that individuals have limited control over occasional contacts with flock mates in this captive environment. I interpret the weak links as the baseline level of interaction. By contrast, I interpret the strong links as the expression of social tactics. In these geese, social tactics would involve preferential bonds with familiar individuals from the gosling phase (Kurvers et al., 2013), which seem to replace the strong bond that normally occurs between parents and offspring and between siblings. I hypothesized that when two bonded individuals interacted with the same unfamiliar individual, this created a weak triplet closed by a strong link. Kurvers et al. raised the goslings in two separate flocks of the same size before merging the two flocks together; therefore, half of the individuals were familiar and half were unfamiliar. Another goose social tactic is to spatially avoid the most dominant, heaviest individuals (Stahl et al., 2001). That avoidance tactic can also generate an excess of weak triplets closed by strong links, typically when two bonded superior competitors both repel a subordinate individual. The original study design afforded the opportunity to compare male and female networks. Although it is a bit tentative, that comparison between sexes might help decipher the two aforementioned proximate mechanisms, because males are more aggressive and exhibit steeper dominance gradients than females.

Throughout, to standardize across datasets and enforce that 0, ≤ *c* ≤ *b* ≤ *a* ≤ 1, I logit-transformed all non-zero weights so that the middle point between the lowest and highest non-zero weights was attributed weight 0.5. Results were qualitatively the same if using a linear transformation.

## Results

### Simulation study

The new method correctly separated the three test scenarios (Fig. 2b: yellow, golden yellow and orange symbols). It not only distinguished the three scenarios from one another, but from other network types as well.

The usual transitivity scores mostly failed at the same task, although their performance depended on the choice of the threshold *c*\*. With no threshold (*c*\* = 0, left panel) or with a high threshold (*c*\* = 0.5, right panel), the usual scores did neither separate the test scenarios from one another, nor from other scenarios, including from scenarios that did not feature any intrinsic excess of one type of triplet. With a moderate threshold (center panel), 𝒞_0_ and 𝒞_1_ separated the three test scenarios by giving them boundary scores 0 or 1. The value of *c*\* to achieve that separation depended on the specifics of the simulation scenarios, here *c*\* = 0.1.

### Illustration 1: grooming networks in rhesus macaque

The kinship structure explained more variation in graph-level transitivity scores than the environment (Fig. 3: triangular symbols). However, the edge density (Fig. 3: color scale) was already capturing that information. The new triad-level scores revealed the extent to which both kinship and captivity influenced the transitive properties: χ^2^3(kin vs. nonkin) = 3793.8; χ^2^3(captive vs. free-ranging) = 842.8 (Fig. 4a-c; both P < 0.001). The difference in transitive properties were much stronger between kin and nonkin groups than between captive and free-ranging groups (Fig. 4a and 4c vs. 4b). The negative quadrant tests (Fig. 4: filled vs. empty histograms) further revealed that the effect of kinship and captivity on the transitive properties operated via the dyadic edge weights, rather than via specific triadic association patterns. Overall, captivity influenced the frequency of weak links and kinship influenced the frequency of strong links.

**Fig. 3:**
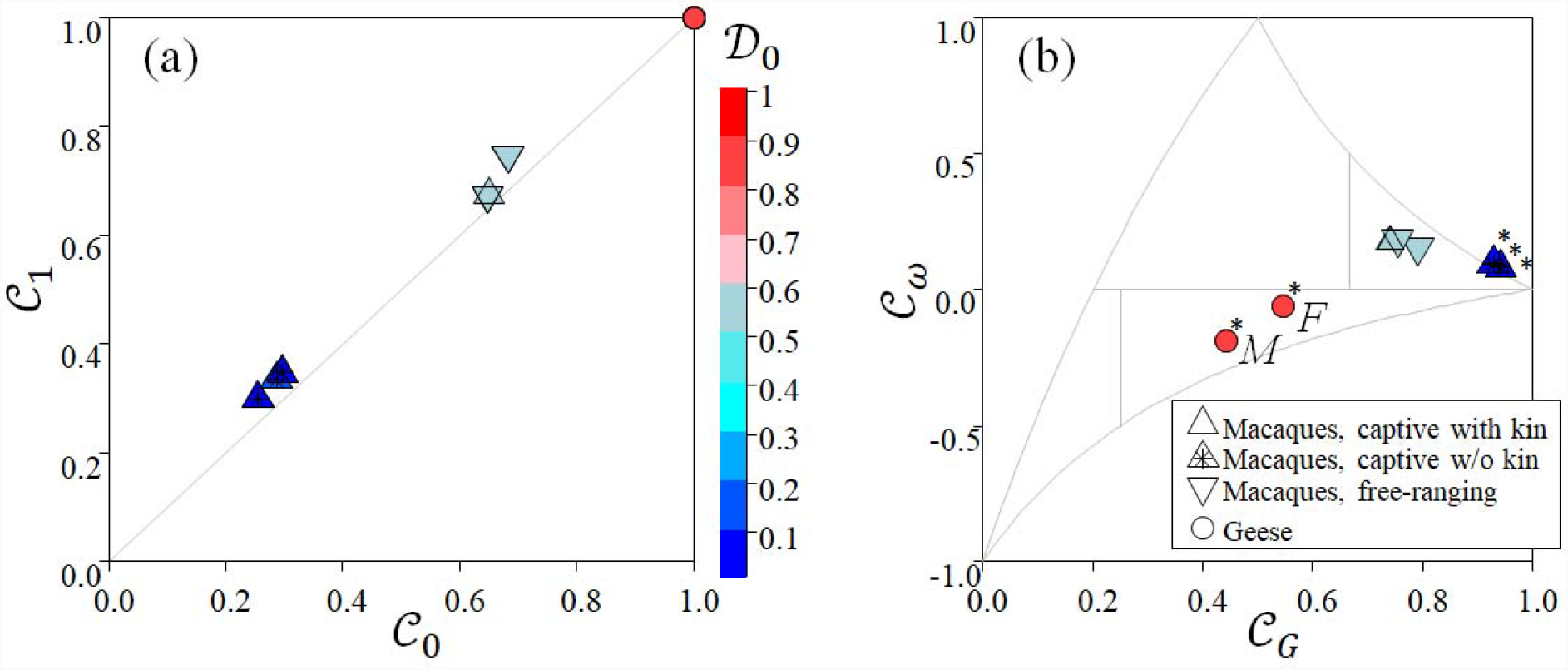
Case studies: graph-level scores. (a) The usual transitivity scores. (b) The new transitivity scores. The asterisks indicate significant distance tests.

**Fig. 4:**
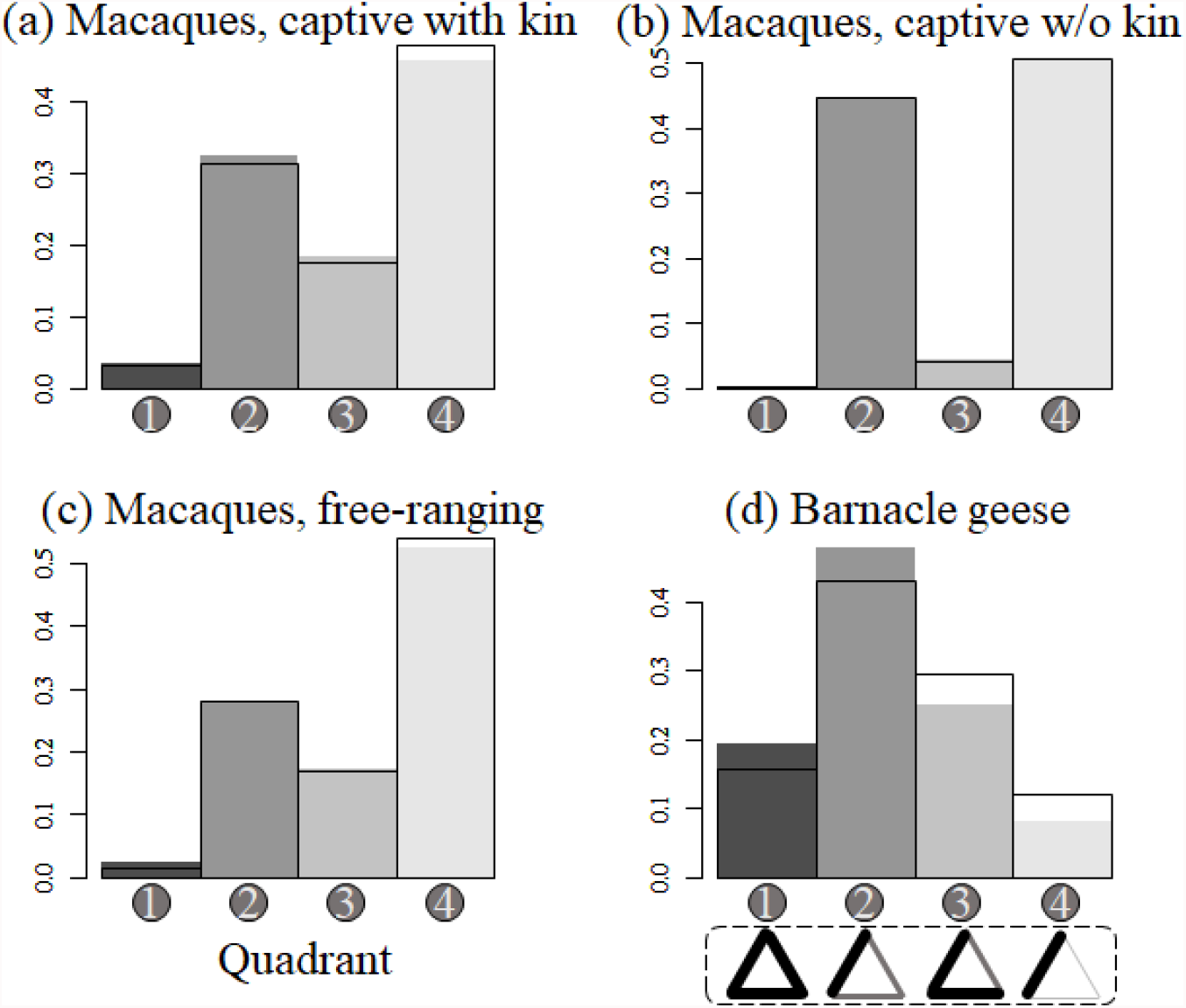
Case studies: triplet-level scores. The triplets are divided into four quadrants as explained in Fig. 1. Grey histograms: observed distributions. Unfilled histograms: expected distributions at random, given the observed distributions of edge weights.

*Illustration 2: barnacle geese experiment*

Since the networks were full, classical metrics failed at uncovering any other information beyond the fact that edge density = 1 (Fig. 3a: red round symbol). By contrast, the new metrics revealed differences between the male and female groups (Fig. 3b: “M” vs. “F”). Both sexes exhibited an excess of weak triplets closed by strong links, as predicted, but the effect was much stronger in males than females. This is consistent with the fact that males are more aggressive and more dominant than females, and suggests a role for the spatial avoidance of dominants in the formation of weak triplets closed by strong links. Contrary to the macaque networks, the quadrant test was very significant for the geese (Fig. 4d: filled vs. empty histograms; P < 0.001). This means that the high frequency of weak triplets closed by a strong link but also the high frequency of balanced triangles (Fig. 4d) did not emerge solely from the dyadic association frequencies. Indeed they resulted from triadic-level processes. This result is strongly consistent with the hypothesis of a triad-based dominance mechanism in which bonded individuals are dominant over singletons.

### Algorithmic considerations

The algorithm in Appendix S1 corresponds to the very most basic option. Its complexity is *O*(*V*^3^), where *V* is the number of vertices in the graph. Even with parallelization, this becomes intractable for *V*>100, especially if using the aforementioned permutation tests. However, I envision some gains in computing time following Schank and Wagner (2005) and references therein. Note however that most animal datasets feature less than 100 individuals.

## Discussion

I expect these weighted network transitivity scores to be most relevant when 1) the networks are dense, meaning that most dyads interact at least once during the study period and 2) the strength of dyadic connections depends on the strength of other connections in the immediate social neighborhood. The barnacle geese case study provided a real-life example of such a situation where the occurrence of a strong dyadic bond suppressed the probability that other strong dyadic bonds would occur in the immediate neighborhood.

The new metrics can also prove useful in large comparative analyses because (1) they capture a different type of information than usual graph summary metrics and (2) they do not require nor depend on the dichotomization of the network using an arbitrary threshold. The appropriate threshold depends on the biological situation (cf. simulation results), making standardization across datasets a bit hazardous. In addition, filtering the statistically insignificant links can be counterproductive for the inference, because the weakest links in a network critically contribute to major processes, such as the spread of epidemics and information from clique to clique (Granovetter, 1973). With the metrics proposed here, there is no need to filter the weakest links. For example, my illustration cases suggest that in captive populations, weak connections represented the baseline level of contact between companions of captivity. The macaque networks were nevertheless still dominated by false or nearly false triplets, as expected due to the high selectivity of grooming interactions. But in the geese, the high baseline rate of contact combined with a few strong bonds made weak triplets closed by strong links the most frequent type of triad (Fig. 4).

Compared to recent trends in network analysis, my relatively ad hoc approach may be perceived as a step back in terms of complexity. Indeed, this is an approach geared towards the study of time-averaged or time-aggregated networks. Modern analysts dealing with big datasets recorded by automated sensors tend to target the instant properties of temporally varying networks (Artime et al., 2017). However, for animal studies, the instant network configuration may not always be particularly relevant, because repeated interactions are required to take effect and/or because the frequency of dyadic contacts is the best summary for functional analysis. Logistically, aggregating data over some span of time also has the advantage of smoothing out any error, missing data, or variation in sampling effort. Such errors are frequent in datasets collected by people from uncooperative animal subjects. They are expected to decrease in frequency with the advent of automated data collection devices, but the latter remain subject to stringent constraints regarding the definition of dyadic contacts, e.g., depending on the detection distance of RFID transponders. In that context, I envision the next step for the study of aggregated contact networks might be to incorporate them within a time-specific capture-recapture framework (Lebreton et al., 2009). This means monitoring the dyadic observation histories that emerge from the presence and detection of individuals, and the occurrence and detection of dyadic interactions between them. A capture-recapture sampling design would in particular allow the explicit modeling of the link between the frequency of contacts and the probably to miss the connection entirely.

## Acknowledgements

The data were procured from the online open source repository ASNR (Sah et al., 2019) (https://github.com/bansallab/asnr). I am very grateful to the authors who made their data available on ASNR, to the creators of the repository, as well as Sebastian Sosa for suggesting it.

## Supplementary material

Appendix S1: R script

